# The roles of *hox 13* genes in newt limb development and regeneration

**DOI:** 10.1101/789180

**Authors:** Takashi Takeuchi, Fumina Minamitani, Kazuki Koriyama, Yukio Satoh, Ken-ichi Suzuki, Shuji Shigenobu, Takeshi Inoue, Kiyokazu Agata, Toshinori Hayashi

## Abstract

Posterior *Hox* genes play crucial roles in limb development and specify regions in the proximal-distal (PD) axis of limbs. However, there is no direct genetic evidence that *Hox* genes are essential for limb regeneration. Moreover, if essential, it is totally unknown which *Hox* genes have the same or distinct functions between development and regeneration. Here, we mutated *hox13* using an efficient CRISPR/Cas9 system in newts (*Pleurodeles waltl*), which have strong regenerative capacities in various tissues. Triple or double mutants of *hox13* paralogs lost their digit and metacarpal/metatarsal bones. Limb regeneration progressed but regenerates lacked the same autopod region. These results showed that *hox13* paralogs have the same functions in limb development and regeneration.

## INTRODUCTION

The questions to what extent regeneration is similar to development, and what events are specific for regeneration have been topics of discussion for a long time. In limbs, several points have been shown to be regeneration specific. For example, signaling from neurons is necessary for blastema growth in limb regeneration (for review see Stocum, 2017) but not for limb bud growth in development. Although previous studies have suggested that pattern formation of limbs is conducted in a similar manner during development and regeneration (for review see Nacu and Tanaka, 2011), it is still unknown whether the same molecular systems are used.

The posterior *Hox* genes play critical roles in pattern formation and also limb growth along proximal-distal (PD) and anterior-posterior (AP) axes during development. These *Hox* functions are involved in determining ZPA and AER activities (for review see Zakany and Duboule, 2007). Analyses of the expression patterns and phenotypes in spontaneous and gene knockout mutations supported a model that *Hox9/10, Hox11* and *Hox12/13* paralogs specify the identity of the stylopod, zeugopod and autopod, respectively, during limb development (Davis et al., 1995; Fromental-Ramain et al., 1996a; Fromental-Ramain et al., 1996b; Wellik and Capecchi, 2003; Zakany and Duboule, 2007). For example, *Hoxa13* and *d13* would specify the distal region in the autopod, because double mutant mice lacking *Hoxa13* and *d13* lost all their digits and metacarpal/tarsal bones (Fromental-Ramain et al., 1996b). In addition, disorders of digit formation, such as hypodactyly, have been linked to human mutations in *HOXA13* and *HOXD13* (for review, (Lappin et al., 2006). However, there is no direct genetic evidence that *Hox* genes are essential for limb regeneration as well as limb development. Furthermore, if essential, it is totally unknown which *Hox* genes have the same or distinct functions between development and regeneration. It is also unknown if the downstream signaling of *Hox* genes is the same or different between development and regeneration. The reason for this is that there had been no way to perform reversed genetics sufficiently in animals which can regenerate their limbs.

Urodele amphibians have the remarkable ability to regenerate many tissues, including limbs. We developed molecular genetic systems in an urodele animal, Iberian ribbed newt (*Pleurodeles waltl*) (Hayashi et al., 2013), and recently established a highly efficient gene knockout system using CRISPR/Cas9 (Suzuki et al., 2018). The system enables us to mutate the target gene in the whole body and almost all alleles (more than 99%) of F0 animals (hereinafter referred as to crispants), due to the longer time for the first cleavage in *P. waltl* (6 h at 25 °C). Therefore, we can disrupt multiple genes in F0 animals by injection of multiple guide RNAs, and analyze the functions without crossing newts. We also performed a comprehensive analysis of a *P. waltl* transcriptome, and established gene models for almost all protein-coding genes in *P. waltl*, including all *hox* genes (Matsunami et al., 2019). In the present paper, using these systems, we analyzed the functions of *hox13* paralogs during limb development and regeneration in newts.

## RESULTS

### *Hox13* crispants could not form their digits during limb development

A comprehensive analysis of *P. waltl* transcriptome revealed that *P. waltl* has at least three *hox13* paralogs (*a13, c13 and d13*) and transcripts of these three genes were identified in the limb blastemas (Matsunami et al., 2019). Because we also found transcripts showing high homology with *hoxb13* of the palmate newt (*Lissortriton helveticusi*, GenBank:DQ158059) and Axolotl (*Ambystoma mexicanum*, GenBank:AF298184.1) in *P. waltl* transcriptome data (Matsunami et al., 2019), we examined mRNA expression of all four *hox13* paralogs (*a13, b13, c13* and *d13*) by RT-PCR in limb blastemas (Fig. 1A). The expression of *hoxa13, c13* and *d13* but not *b13* was detected (Fig. 1A). The data is consistent with that in *P. waltl* transcriptome data (Matsunami et al., 2019). Therefore, we decided to disrupt *hox a13, c13* and *d13* using a CRISPR/Cas9 system.

**Figure 1.**
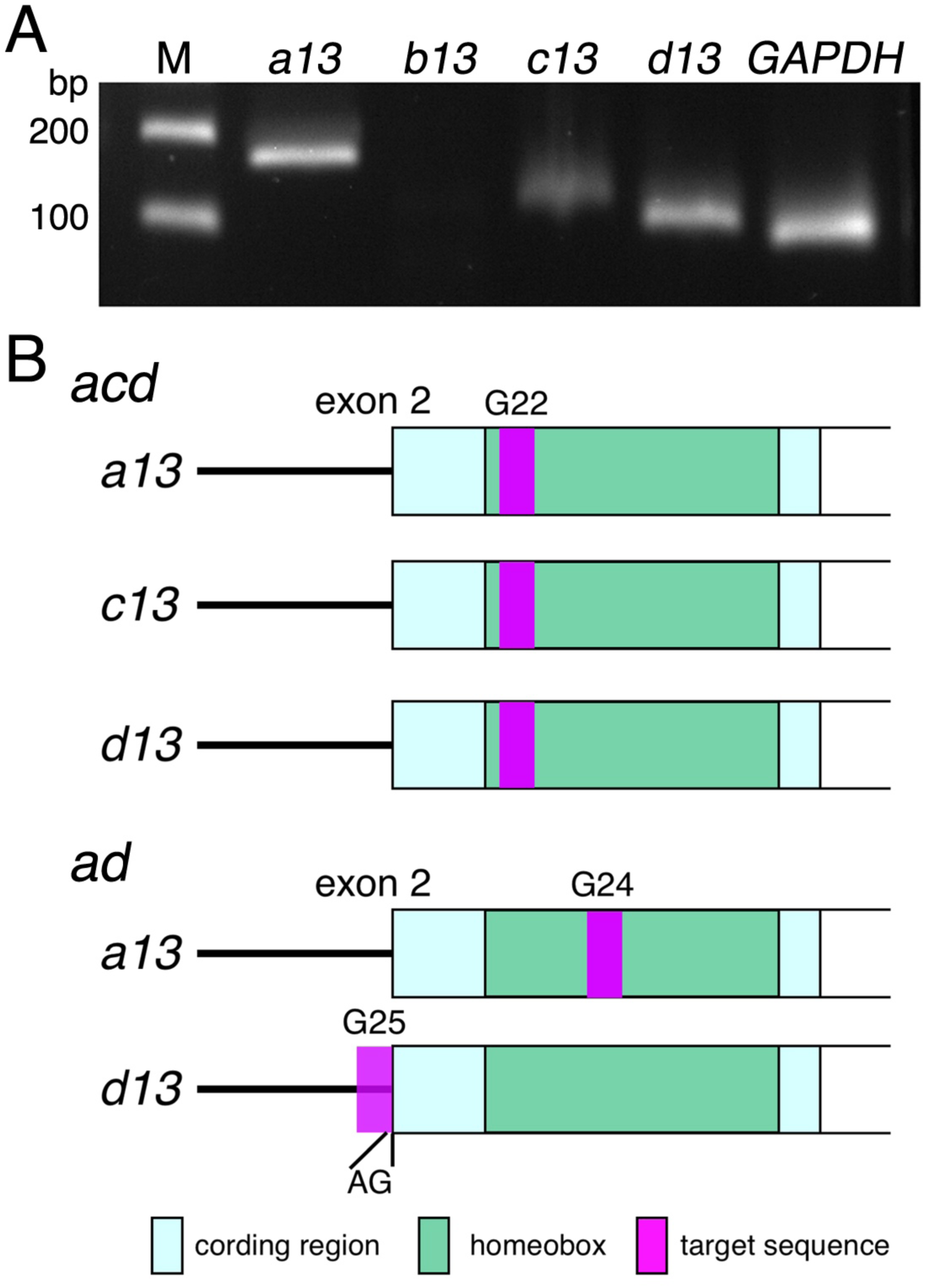
Expression of *hox13* paralogs and positions of target sequences for CRISPR/Cas 9 mediated gene disruption. (A) Expression of *hox13* paralogs in the late blastemas was analyzed by RT-PCR. M: DNA markers (100 and 200 bp). *GAPDH* was used as a positive control. (B) Schematic representation of target sites in *hox13* genes. PAM sequences were not included. AG in *ad* indicated consensus AG sequence in 3’ splice site of the first intron of *hoxd13*.

We used two sets of guide RNAs; *acd* (G22) and *ad* (G24 and G25) (Fig. 1B). G22, G24 and G25 were the names of the CRISPR RNAs (crRNAs). *acd* and *ad* were designed to mutate three paralogs (*a13, c13* and *d13*) and two paralogs (*a13* and *d13*), respectively. G22 had complete homologies with one identical sequence in the homeoboxes of *hoxa13* and *d13* and high homology with a sequence in *c13* (19 matched nucleotides among a total of 20 nucleotides of the protospacer sequence). G24 and G25 had complete homology with another sequence in the homeobox of *hoxa13* and with a sequence around the splicing acceptor of exon 2 in *hoxd13*, respectively. Disruption of *c13* by G24 was maybe possible, because it had slightly high homology with a sequence in *c13* (18 matched nucleotides among a total of 20 nucleotides of the protospacer sequence). All target sequences had a protospacer adjacent motif (PAM) sequence. All three crRNAs showed no high homology (more than 3 nucleotides were unmatched among 20 nucleotides of the protospacer sequences which further had PAM sequences at the 5’ or 3’ end) with all other gene bodies in *P. waltl* comprehensive transcriptome data, including the sequences of other Homeobox genes (Matsunami et al., 2019).

The ribonucleoprotein complexes (RNPs) including Cas9 and each guide RNA set were injected into fertilized eggs. Uninjected siblings (wild type animals, N=28) showed normal limb development (Fig. 2 and 3). All 9 animals to which *acd* guide RNA was injected (*acd* crispants) and 15 among 18 *ad* crispants lacked all fore-limb and hind-limb digits (Fig. 2A). The remaining three animals in *ad* crispants had only 1, 2 and 4 digits in their four limbs, respectively, indicating that they lacked 14-17 digits. Bone staining of limbs without digits showed that the stylopod and zeugopod bones formed and appeared to be normal, however, these limbs had neither phalanges nor metacarpal/metatarsal bones (Fig. 2B). The morphology and number of carpal/tarsal bones were abnormal (Fig. 2B). Some carpal/tarsal bones were sharp and the tip stuck out from the skin (Fig. 2B, arrowheads). Observation of limb development in crispants showed the elongation of limb buds, but no digits were formed (Fig. 3). The phenotypes (loss of phalanges and metacarpal/metatarsal bones) are consistent with those in *Hoxa13* and *d13* double mutant mice (Fromental-Ramain et al., 1996b).

**Figure 2.**
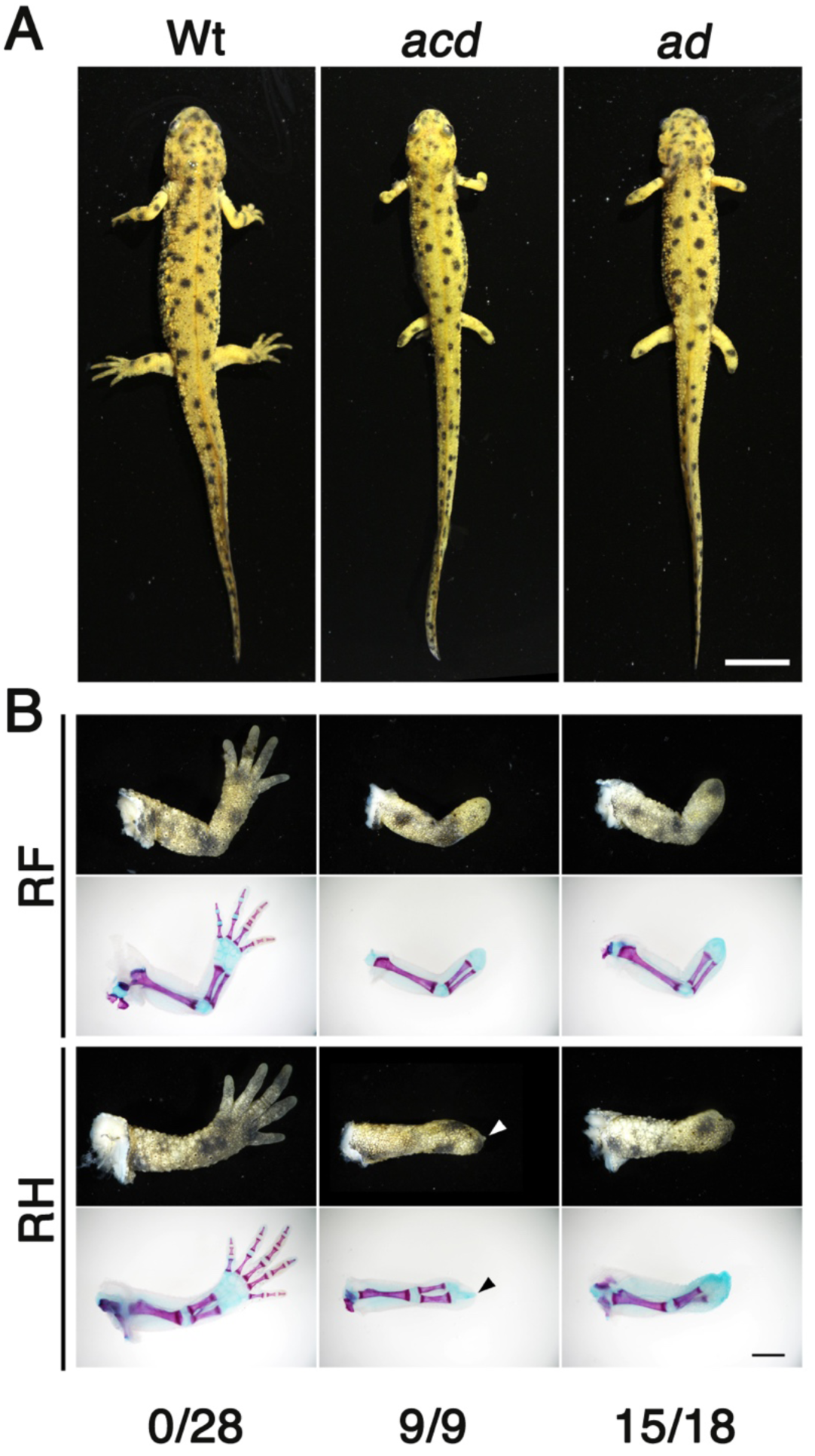
*Hox13*crispants lacked their digits. (A) Dorsal whole mount views of representative wild type (Wt), *acd* and *ad* animals at 16 weeks post fertilization (wpf). No digits were formed in *acd* and *ad* crispants. Bar, 1 cm. (B) Bone staining patterns of right forelimbs (RF) and hindlimbs (RH) of representative wild type (Wt), *acd* and *ad* animals at 16 wpf. Anterior, up. Limbs of *acd* and *ad* animals had neither phalanges nor metacarpal/metatarsal bones. Arrowheads show an example of sharp carpal/tarsal bones. Numbers shown below indicate numbers of animals with no digits / numbers of total animals. Bar, 2 mm.

**Figure 3.**
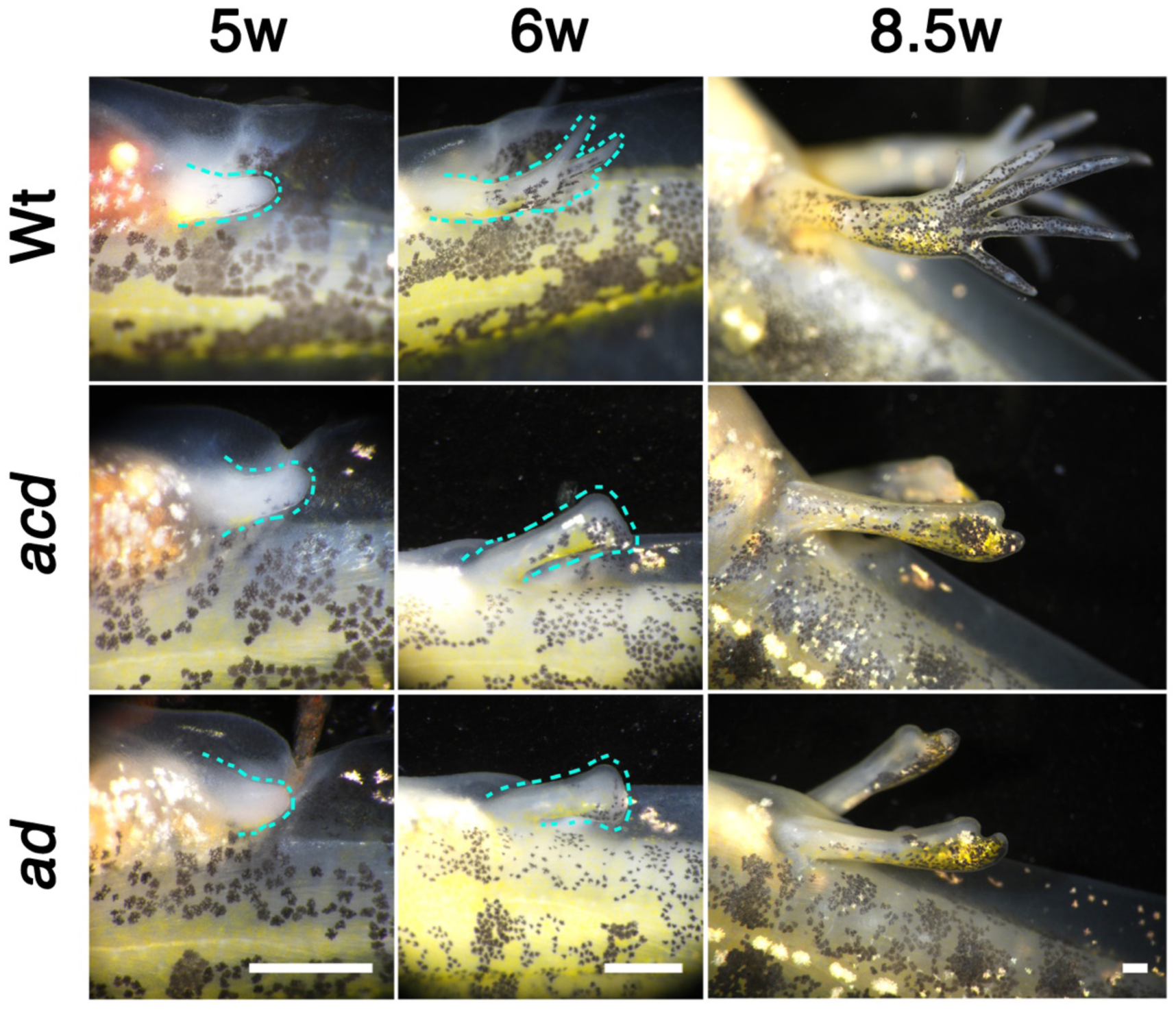
Limb buds elongated but did not form any digits during limb development. Development of right hindlimb. Limbs at 5, 6 and 8.5 wpf are shown. Ventral, up and anterior, left. Outlines of limb buds are shown by light blue dotted lines. Bar, 1 mm.

### *Hox13* genes were mutated in the crispants

Because the target sequences of G22 in *acd* crispants had a cleavage site for PvuII restriction enzyme, we examined the digestion of amplicons of *hoxa13, c13* and *d13* in *acd* crispants to analyze somatic mutations (Fig. S1A). Insufficient or no PCR products were obtained in some cases or the sizes of the main or extra products were smaller than that of a wild type animal, suggesting deletion. Similar results were obtained in the cases of *ad* crispants (Fig. S1B). For most products from *acd* crispants in all three genes, which were well amplified, digestion was not detected (Fig. S1A), suggesting mutations in the PvuII site. Some products were digested partially. Since mutations can occur in nucleotides at sites other than a PvuII site, we could not conclude that the digested fragments resulted from wild type alleles.

We also performed amplicon sequencing by next generation sequencing (NGS) to conduct a detailed analysis. Genomic DNAs of *hoxa13, c13* and *d13* in both *acd* and *ad* were investigated. Two crispants of each set, which lost all digits, were examined. *Hoxc13* was mutated in *acd* crispants (mutation rates, 89.2 and 99.8%), but not in *ad* crispants. Almost all alleles (99.7-100%) in *hoxa13* and *d13* were mutated in all analyzed animals. One to 4 mutant alleles occupied over 96% of all alleles examined (Table 1, 2, S1 and S2). Mutations in homeoboxes (*a13, c13* and *d13* in *acd* and *a13* in *ad*) yielded frameshift mutations or deletions of 2-5 amino acid residues within K10-L16 in helix I (Fig. S2) including conserved residues among homeodomains of posterior Hox proteins (Fig. S2). Helix I is critical for homodimerization, which is required to activate transcription (Zhang et al., 2011). These frameshift mutations and deletions suggested the loss of function of *hox13* genes (see also Discussion).

**Table 1.**
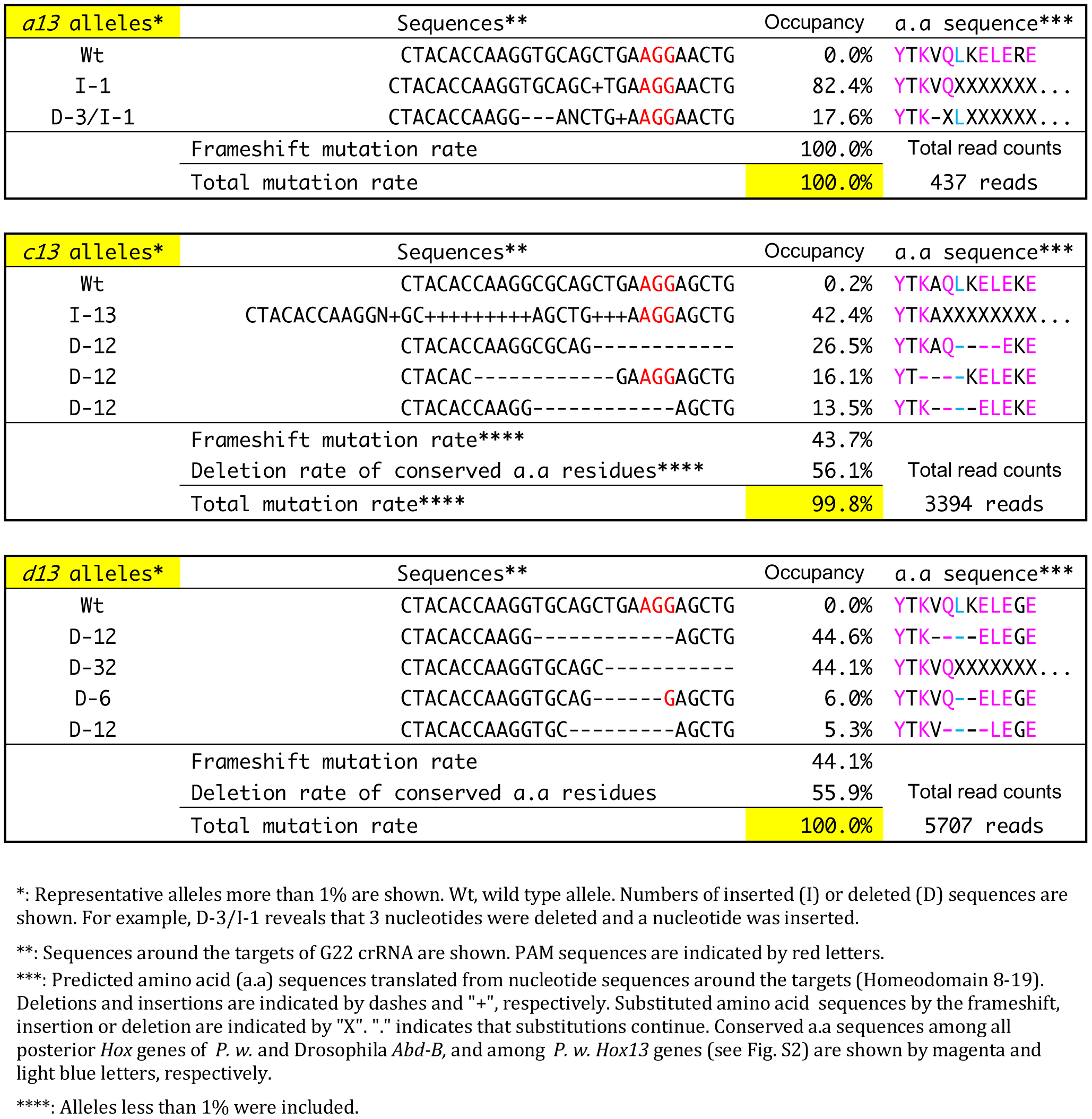
Genotypes of a representative *acd* crispants (#5) analyzed by amplicon sequencing

**Table 2.**
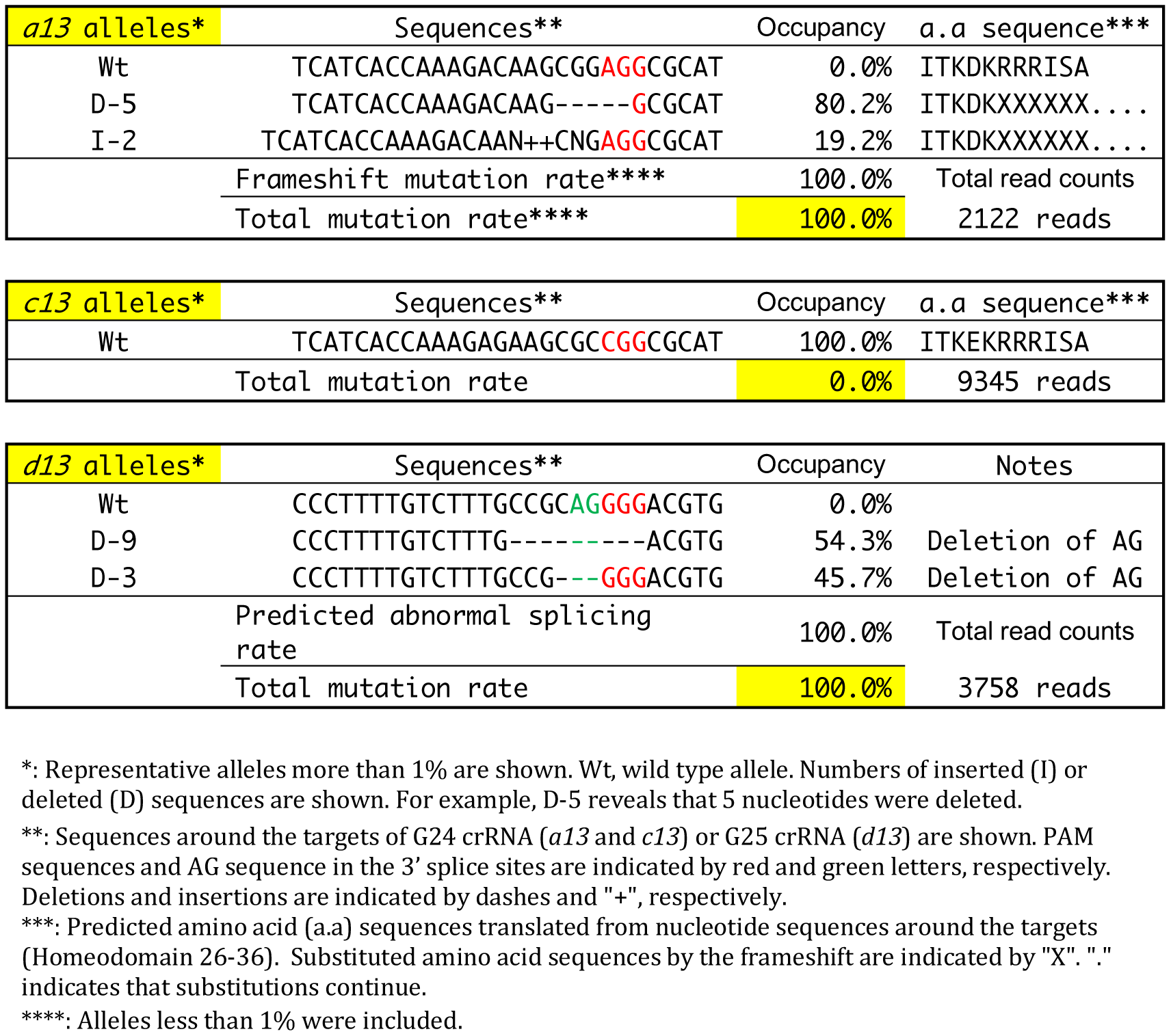
Genotypes of a repelsentative *ad* crispant (#16) analyzed by amplicon sequencing

Deletions or insertions around the splicing acceptor of *hoxd13* in *ad* (Table 2 and Table S2) also suggested loss of function of *hoxd13* by abnormal splicing. In particular, deletion of the consensus AG sequence in the 3’ splice site makes it impossible to splice at the normal site. Even if new potential splice sites work, the splicing yields frameshift mutations or large deletions in the homeodomain.

One animal in *ad* (*ad*#16) showed frameshift mutations and deletion of AG sequence in the 3’ splice sites in all alleles of *hoxa13* and *hoxd13*, respectively (Table 2). Therefore, both the *hoxa13* and *d13* genes were almost certainly disrupted in at least this animal. All newts in *acd* and *ad* including #16 showed the same phenotype in the autopod (loss of phalanges and metacarpal/metatarsals), which was also observed in *Hoxa13* and *d13* double knockout mice, suggesting that at least *hoxa13* and *d13* were disrupted in *hox13* crispants.

### Limb regeneration progressed but regenerates lacked digits in *hox13* crispants

*Hox 13* crispants grew and metamorphosed, although 5 among 9 animals in *acd* and 7 among 18 animals in *ad* died mainly around the time of metamorphosis. One possible reason was difficulty in swimming up to the surface of the water to breath without digits. However, the fact that almost half of the animals survived after metamorphosis was very important for the analysis of limb regeneration, and was different from *Hoxa13* and *d13* double mutant mice which showed embryonic lethality. The fore-limbs of young crispants after metamorphosis (after 16 weeks post fertilization), which had lost all digits, in *acd* (4 animals) and *ad* (10 animals) and uninjected siblings (wild type, 11 animals) were amputated at the proximal region in the stylopod. All four animals in which *hox13* mutations were identified by amplicon sequencing (Table 1, 2, S1 and S2) were included. Regenerates of all wild type animals showed normal morphology (Fig. 4, 5 and movie 1). During limb regeneration, no apparent phenotypes were observed before the notch stage, however, no crispants showed notch structure or digit formation (Fig. 4 and movie 2). Bone staining of limbs showed that all regenerates lacked all digits and metacarpal/tarsal bones in the limb regenerates. The bone structure of regenerates was very similar to that of developed limbs before amputation (Fig. 5). These results first showed that *hox13* has the same function (patterning of the autopod) in limb regeneration as well as limb development. We could not find any regeneration specific phenotypes.

**Figure 4.**
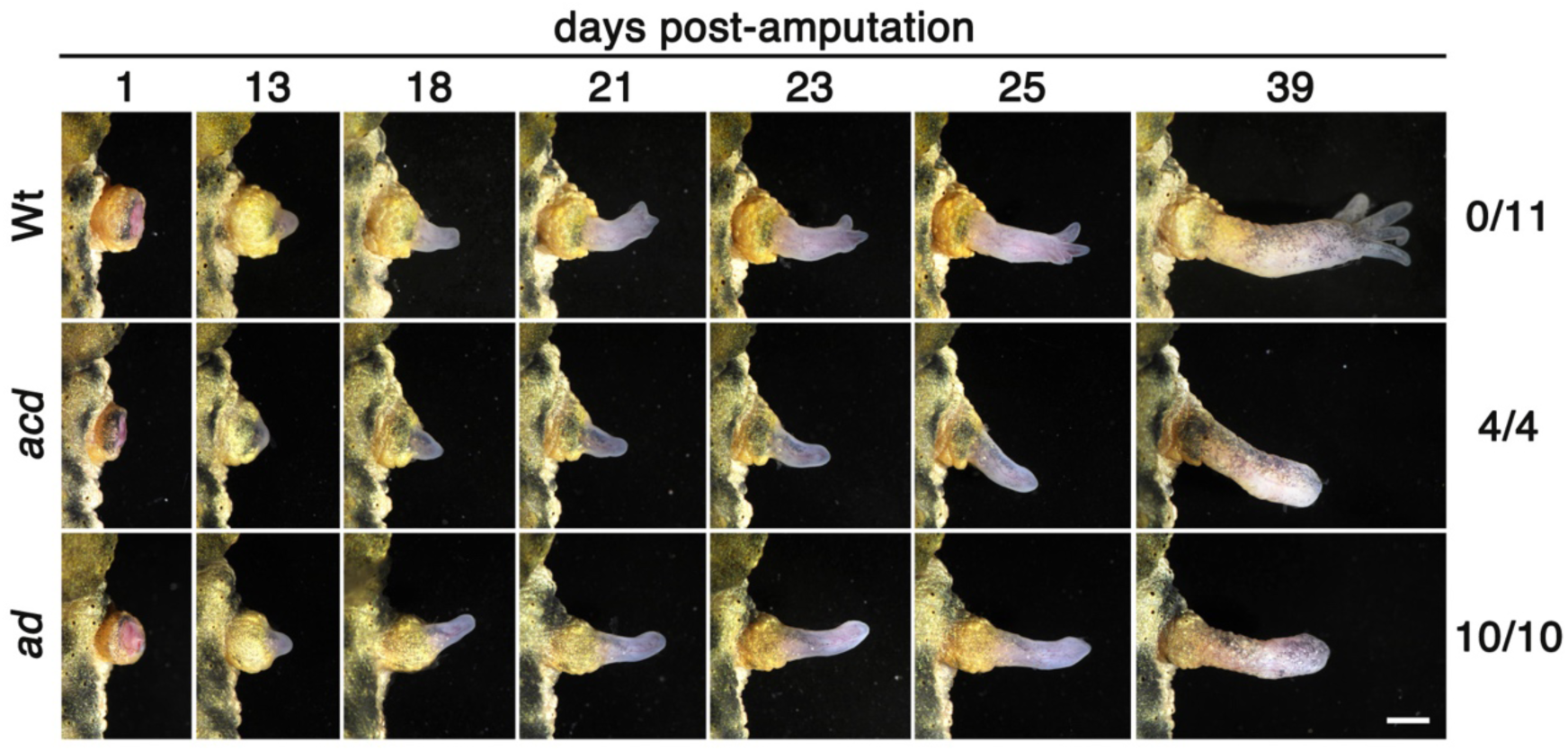
Limb regenerates of *hox13*crispants lacked digits. Dorsal views of representative regenerates of wild type (Wt), *acd* and *ad* animals. Limbs were amputated at 25 weeks post fertilization (over 5 months post fertilization) at the most proximal region in the stylopod. Days post-amputation are shown above. Numbers shown on the right side indicate numbers of animals with no digits / numbers of total animals. Anterior, up. Bar, 2 mm.

**Figure 5.**
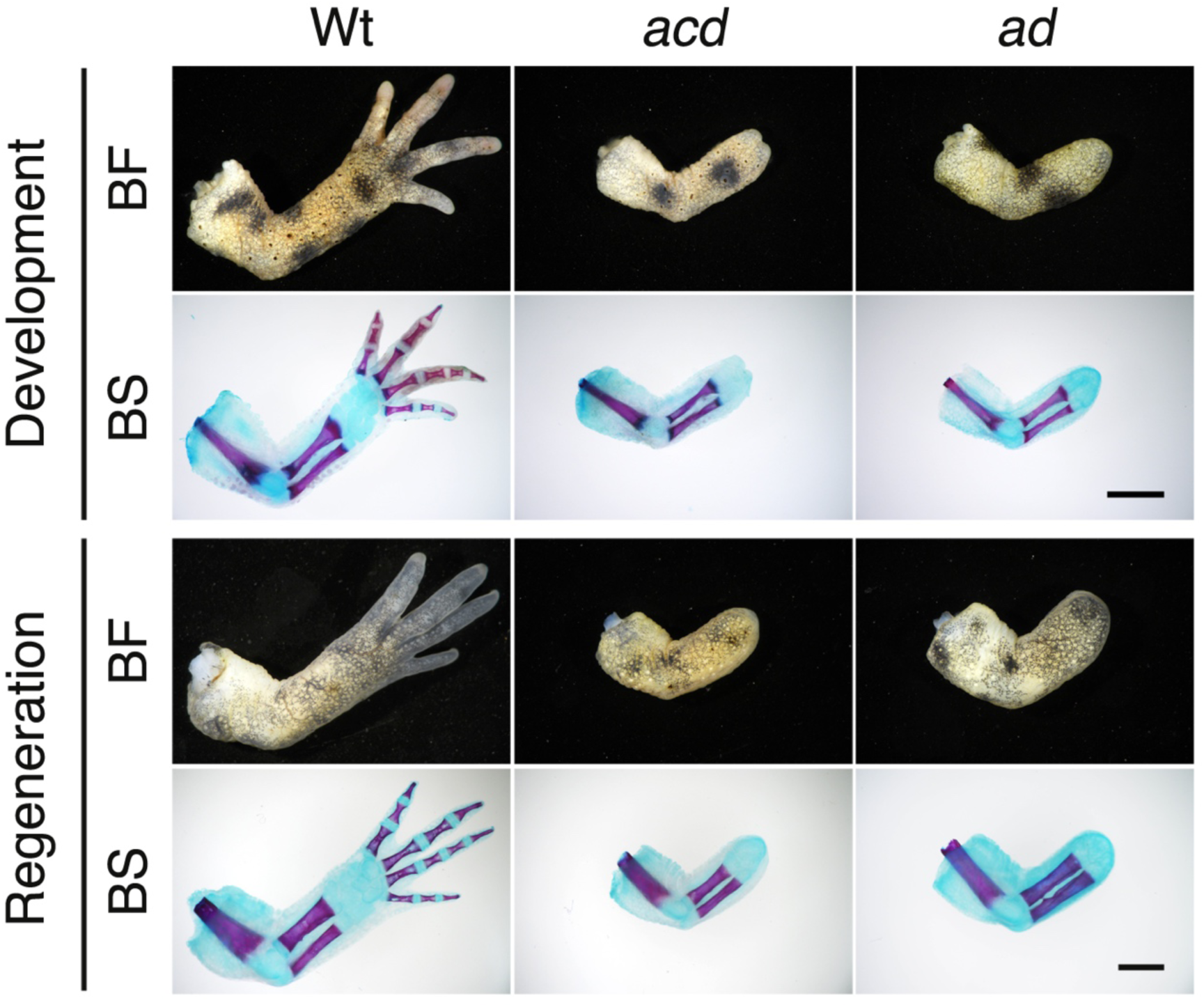
*Hox13* has same function in limb development and regeneration. Dorsal views of representative developed (upper panels) and regenerated (lower panels) right forelimbs of wild type (Wt), *acd* and *ad* animals. Bright field (BF) and bone staining (BS) patterns are shown. For regeneration, limbs were amputated at the most proximal region in the stylopod, and regenerates were fixed at 9 weeks post-amputation. Anterior, up. Bar, 2 mm.

## DISCUSSION

Our results showed that crispants, in which *hox13* genes were mutated, lacked the distal autopod region of the limbs during development and regeneration, suggesting that *hox13* was essential for morphogenesis of the autopod region in newts as well as mammals, and in both development and regeneration.

The results presented in this paper suggest that *hox13* genes were disrupted. Frameshift mutations cause the substitution of downstream amino acid residues and/or truncation by abnormal stop codons, resulting in the loss of over 50% of homeodomains. The lost regions include the recognition helix (helix III, Fig. S2), which is essential for Hox functions. Deletions in helix I would also affect Hox function. Two residues in helix I of HOXA13 (corresponding residues are R18 and E19 in Fig. S2) are critical for dimerization, which is required to activate transcription (Zhang et al., 2011). Deletions of 2-5 amino acid residues within K10-L16 of helix I (Fig. S2), close to the critical residues for dimerization (R18 and E19), would affect the homodimerization by truncation or structural abnormalities in helix I. Therefore, mutations observed in homeoboxes suggest the loss of function of *hox13* genes.

*acd* and *ad* animals exhibited the same phenotypes in limbs (loss of digits) (Fig. 2-5). These data showed that mutations in at least *hoxa13* and *d13* caused the phenotypes. We could not find any *hoxc13* specific functions in the limb development and regeneration. However, it is still possible that disruption of *hoxc13* showed specific or similar phenotypes and that these were masked by a deficiency of *hoxa13* and *d13.* Analysis of single mutants of *hoxc13* in newts should be able to examine this possibility.

We believe that the phenotypes shown here did not result from off-target effects which mutated genes other than the *hox13* gene, for the following reasons. (1) Target sequences were carefully determined, and the sequences used did not show high homology with any other gene bodies in *P. waltl* comprehensive transcriptome data (more than 3 mismatches). (2) Sequences with more than two mismatches would not be mutated, because *hoxc13* was not mutated by G24. (3) All crispants in *acd* and *ad* showed the same phenotypes (loss of digits), although different crRNAs were used. It is almost impossible that another gene was disrupted both in *acd* and *ad*, or that different genes which showed the same phenotypes were disrupted in each Set.

Evolutionally, amphibians acquired limbs with the autopods found in present-day tetrapods. How did amphibians acquire the autopods? *Hox13* genes have been considered to play important roles in acquisition of the autopod structure, and changes in patterns and activities of *Hox13* genes may be involved in acquisition (for reviews see (Leite-Castro et al., 2016; Paço and Freitas, 2018; Schneider and Shubin, 2013; Tanaka, 2016; Woltering and Duboule, 2010). Loss of function experiments, using the CRISPR/Cas9 system, on *hoxa13* and *d13* in zebrafish showed a marked reduction of fin ray, suggesting that digits originated via the transition of the distal structure of the fin (Freitas et al., 2012; Nakamura et al., 2016). As described above, *Hoxa13* and *d13* would specify the distal region in the autopod of mice (Fromental-Ramain et al., 1996b). Our studies shown here connect the mouse to zebrafish in terms of *Hox13* functions, and indicate that *Hox13* genes are essential for formation of the distal region of the appendages (fins and limbs) in mammals, amphibians and fishes. Studies on expression patterns and cis elements of *hox13* genes in newts, and comparisons with those of mice and fish will be important. Another fundamental question is what is the biological significance of the acquisition of digits? If we can use animals without digits for physical and/or behavioral experiments, we should be able to investigate the functions of digits and their significance in evolution. It is difficult to use *Hox13* mutant mice due to embryonic lethality. However, the newt mutants shown here can be used for these experiments. These studies will provide important insight for understanding the molecular mechanisms that enabled the fin to limb transition, and its biological significance.

## MATERIALS AND METHODS

### Newts

In this study, we used Iberian ribbed newts (*Pleurodeles waltl*) which were raised in our laboratory. The animals were reared as described previously (Hayashi et al., 2013). All procedures were carried out in accordance with the Institutional Animal Care and Use Committee of Tottori University (Tottori, Japan) and national guidelines of the Ministry of Education, Culture, Sports, Science & Technology of Japan.

### RT-PCR

Four blastemas at late bud (Koriyama et al., 2018) were used for sampling of mRNA. RT-PCR was performed as described in a previous study (Toyoda et al., 2003). Primers are shown in Table S3. *GAPDH* was used as the endogenous control.

### Preparation of RNPs and microinjection

crRNAs were designed by using CRISPR-direct (Naito et al., 2015). Positions of the targets are shown in Fig. 3 and the sequences are G22: CTACACCAAGGTGCAGCTGA<u>AGG</u>, G24: TCATCACCAAAGACAAGCGG<u>AGG</u> and G25: CCCTTTTGTCTTTGCCGCAG<u>GGG</u> (PAM sequences are undelined). The target sequence of G22 has PvuII cleavage site (CAGCTG). The synthetic tracrRNA, crRNA and Cas9 protein were obtained from Integrated DNA Technologies (IDT; Skokie, IA, USA). The tracrRNA and crRNA were annealed, and then RNPs including Cas 9 were produced in accordance with the manufacturer’s instructions just before injection. Microinjection of RNPs was performed based on our previous reports (Hayashi et al., 2019; Suzuki et al., 2018).

### Limb amputation and bone staining

Limb amputation and bone staining were performed as described previously (Koriyama et al., 2018). The fore- and hind-limbs were amputated at the most proximal region in the stylopod. The developed and regenerated limbs were fixed in 10% formalin-Holtfreter’s solution at 4°C for more than 2 days and stained with Alizarin Red and Alcian Blue.

### Genotyping

Genomic DNA was extracted from tail fins. The target region was amplified from the lysates using EX-Taq enzyme (for conventional PCR analysis, Takara Bio, Japan) or KAPA HiFi enzyme (for NGS analysis, Roche Diagnostics, Basel, Switzerland) with primer sets (Table S3). Digestion of amplicons of *acd* crispants was examined using PvuII (New England BioLabs Japan, Japan). Amplicons for NGS analysis were obtained by primers containing barcoded overhang adaptor sequences according to a 16S Metagenomic Sequencing Library Preparation Kit, and purified using AMPure XP (Beckman Coulter, Pasadena, CA, USA). All amplicons were mixed and subjected to an Illumina MiSeq run (paired-end 300; Macrogen). Sequencing data was analyzed according to our previous study (Suzuki et al., 2018).

### Observation of developing and regenerating limbs

The developing and regenerating limbs were observed with a stereoscopic microscope (Leica, MZFL III) and images including limbs before and after bone staining were acquired with a microscopy camera (Nikon, DS-Ri2).

## Acknowledgements

This work was supported by NIBB Collaborative Research Program (18–204) to T.H.

## Supplemental Figures, tables and movies

**Fig S1.**
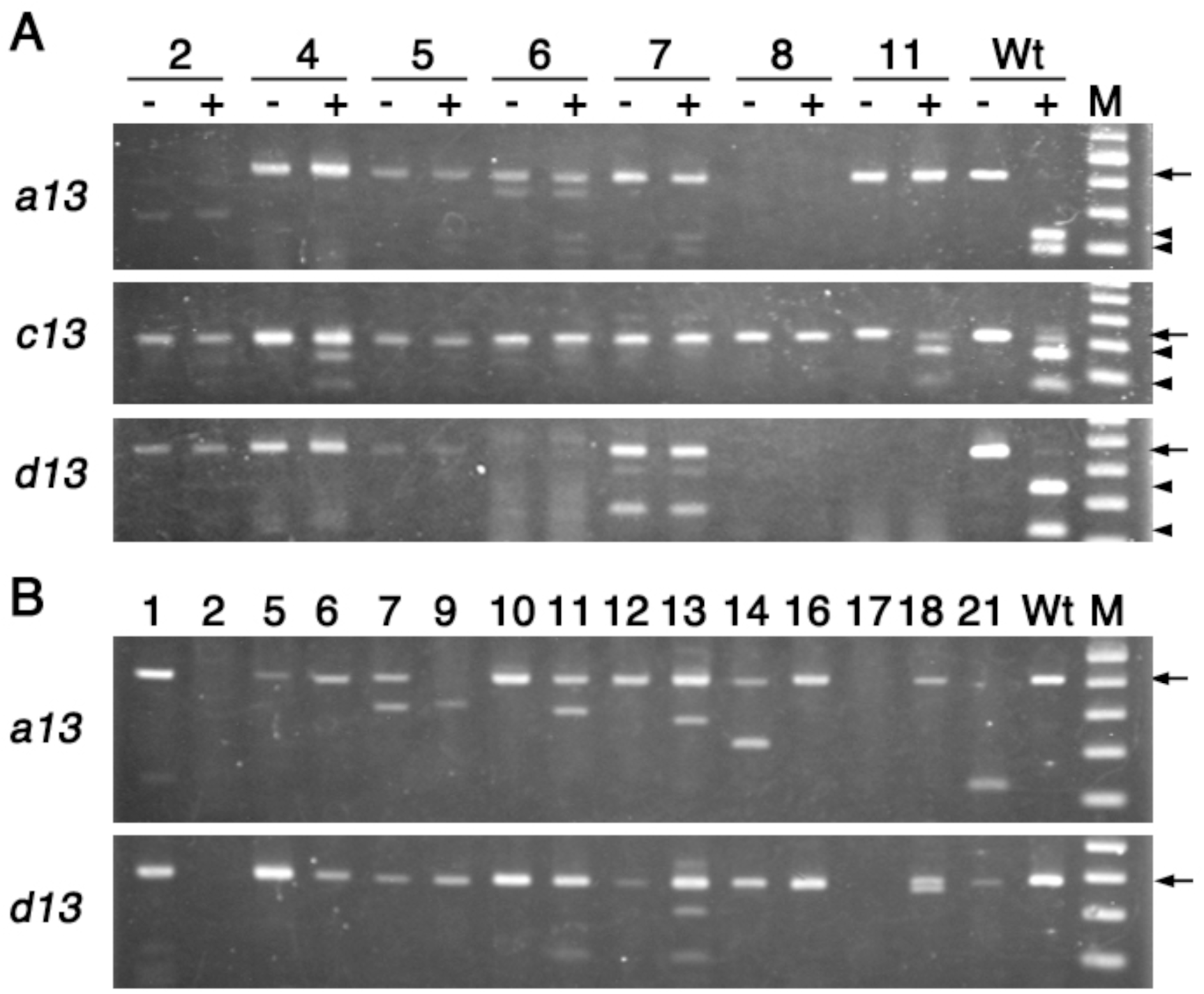
Genotype analysis using amplicons. (A and B) Wt and M indicate data from a wild type animal and size marker, respectively. (A) Genotypes of *acd* crispants were analyzed using restriction enzyme for amplicons. Digestion by PvuII was examined in PCR products of *acd* crispants. Data of representative 7 animals is shown. Numbers above gel photos represent individual identification numbers. “-” and “+” indicate no treatment and treatment with PvuII, respectively. Arrows and arrowheads show positions of undigested and digested amplicons, respectively. (B) Amplicons of *ad* crispants were analyzed by gel electrophoresis. Data of representative 15 animals is shown. Numbers above gel photos represent individual identification numbers. Arrows indicate positions of an amplicon from a wild type animal.

**Fig S2.**
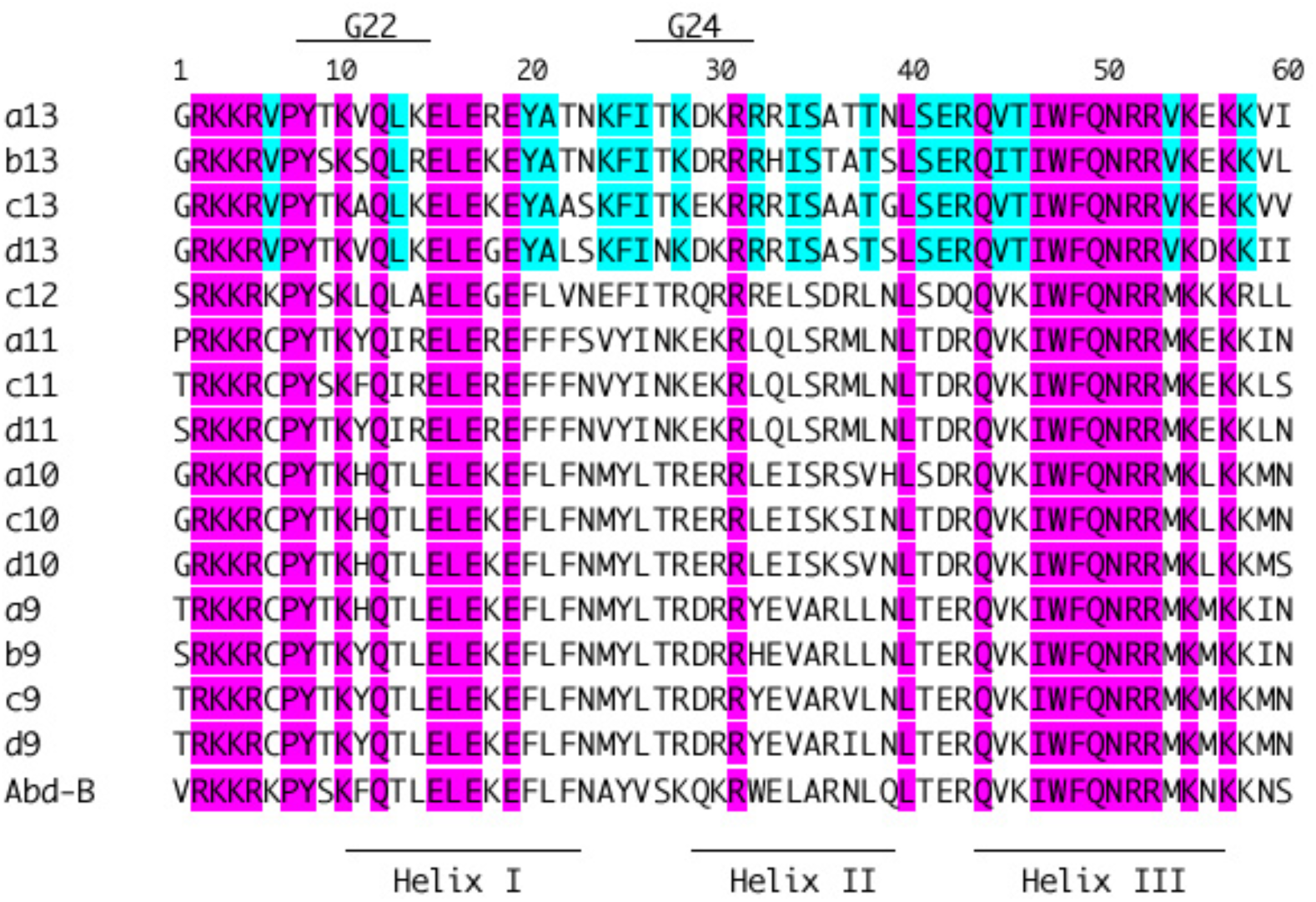
Conserved amino acid residues in homeodomains among *P. waltl* posterior Hox proteins and *Drosophila melanogaster* Abd-B. Magenta and light blue characters represent conserved residues among *P. waltl* posterior Hox proteins and *Drosophila melanogaster* Abd-B and among *P. waltl* Hox13 proteins, respectively. The positions of crRNA (G22 and G23) and helixes are indicated above and below the sequences, respectively.

**Table S1.**
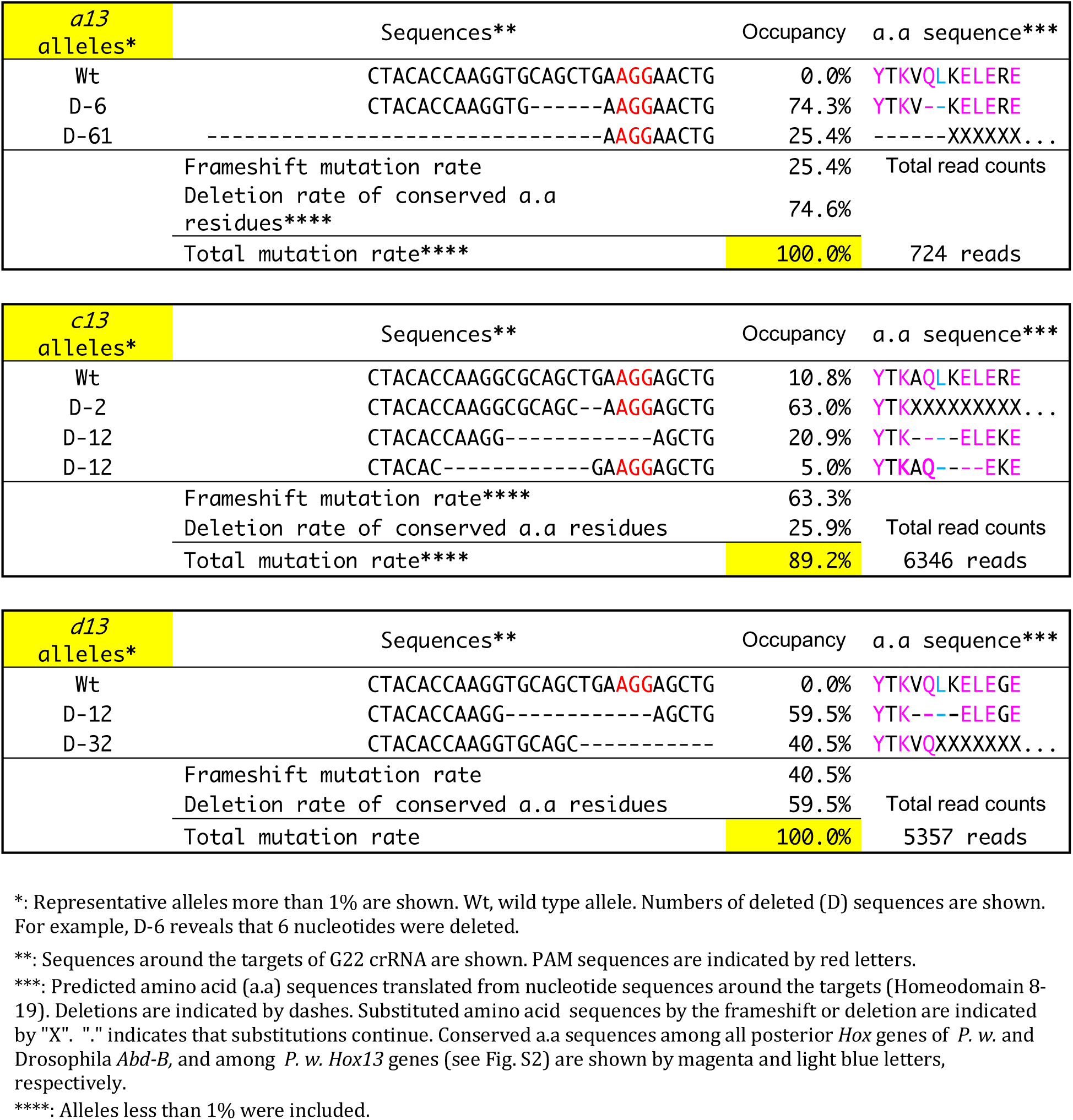
Genotypes of another *acd* crispants (#6) analyzed by amplicon sequencing

**Table S2.**
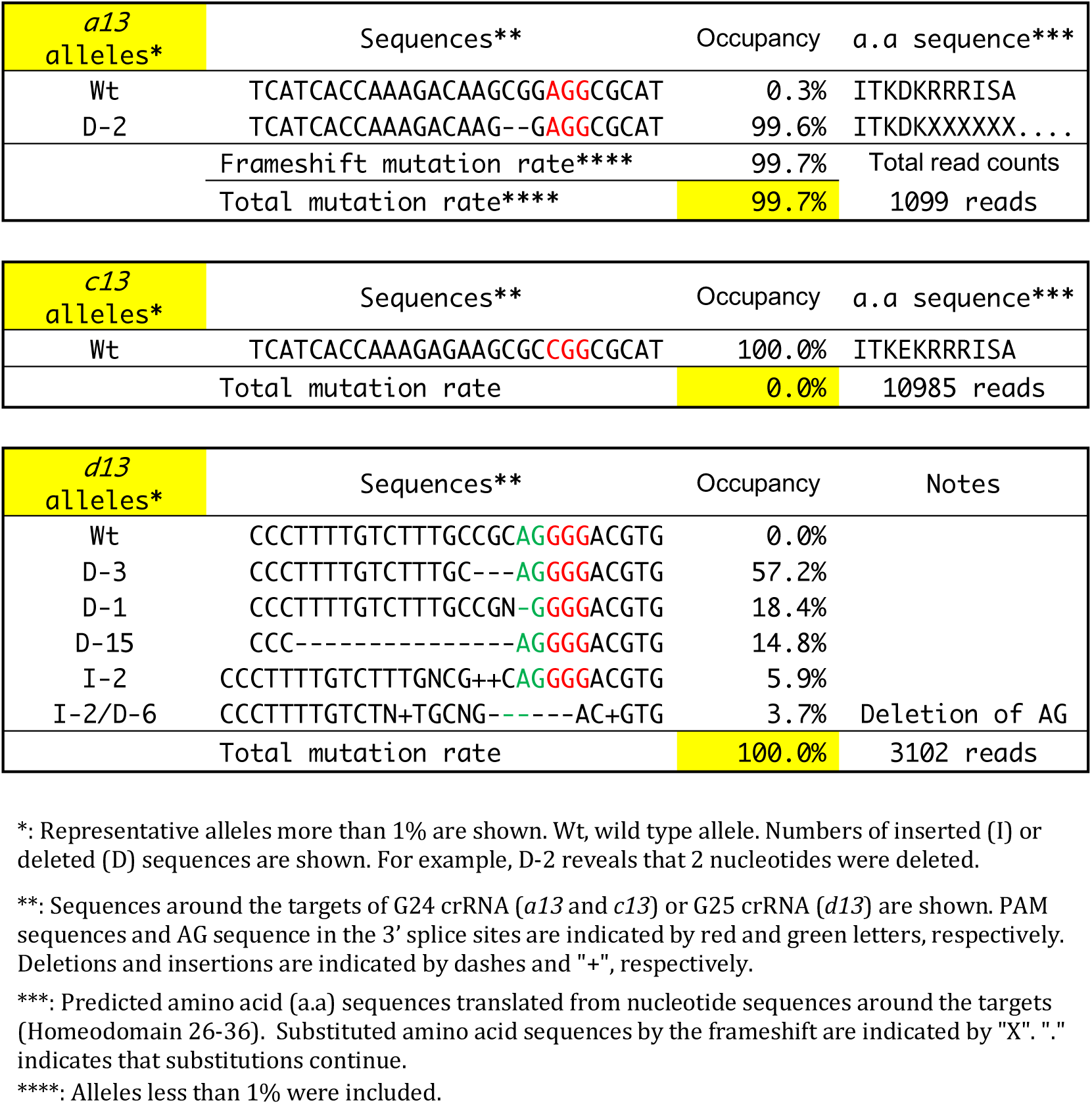
Genotypes of another *ad* crispant (#1) analyzed by amplicon sequencing

**Table S3.**
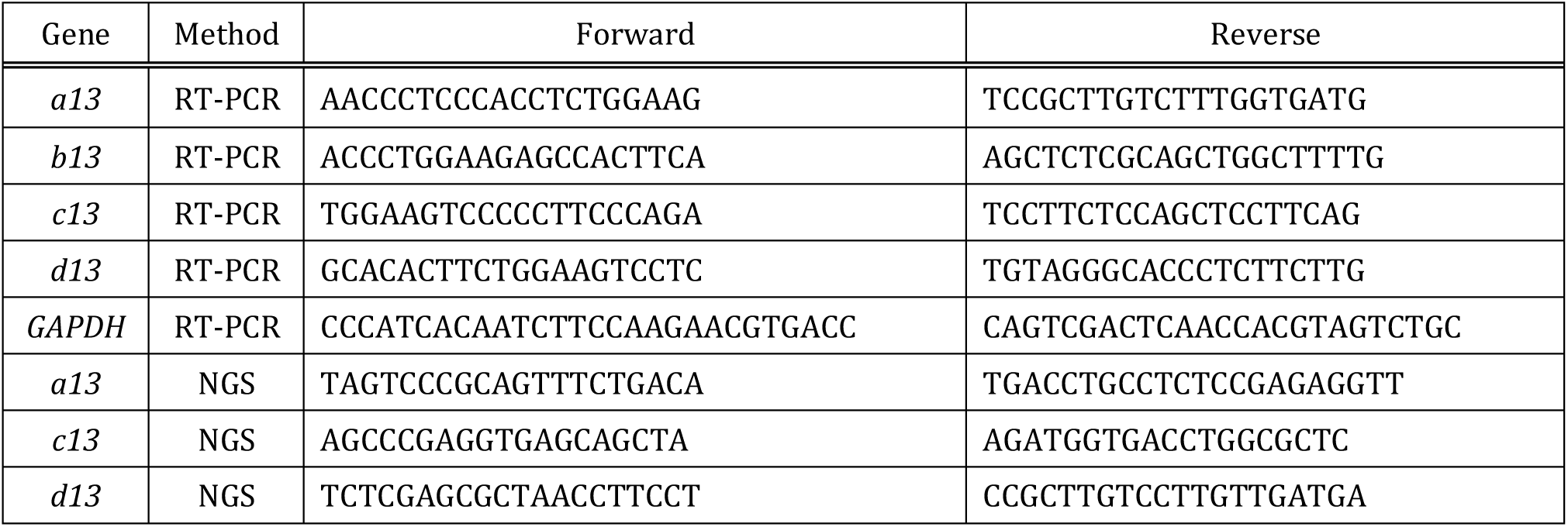
Sequences of PCR primers

**Movie 1. Regeneration of the right forelimb of a wild-type animal**

**Movie 2. Regeneration of the right forelimb of *ad* animal**

